# PIMENTA: PIpeline for MEtabarcoding through Nanopore Technology used for Authentication

**DOI:** 10.1101/2024.02.14.580249

**Authors:** Valerie van der Vorst, Marijke Thijssen, Bas J. Fronen, Arjen de Groot, Margot A.M. Maathuis, Els Nijhuis, Marcel Polling, Joost Stassen, Marleen M. Voorhuijzen-Harink, Robbert Jak

## Abstract

DNA metabarcoding has become a cost-effective method to assess species composition of mixed samples. Developments such as advances in sequencing technology and increased species coverage of reference databases can be leveraged to gain more insights from metabarcoding experiments, given suitable tools. To this end, we introduce PIMENTA, a new pipeline that streamlines the analysis of Nanopore DNA metabarcoding sequencing data.

PIMENTA consists of four phases: pre-processing, clustering per sample, reclustering of all samples, and taxonomic identification. PIMENTA expands a workflow created by Voorhuijzen-Harink et al. Multiple updates have been made, including parallelization of the analysis of multiple samples with the use of high-performance computing (HPC), implementation of a local taxonomy database, and expansion of the taxonomic results summary. Settings have been optimized to process higher quality nanopore reads, for an increased accuracy of taxonomic identification.

We evaluated the pipeline with mock samples of zooplankton species, incorporating COI, 18SV4, and 18SV9 marker sequences. The performance and runtime have been benchmarked against two other existing pipelines. PIMENTA was able to quickly identify species with a high resolution and minimal misidentifications.

## 1. Introduction

In recent years, DNA metabarcoding has become a common technique for the cost-effective assessment of species composition in mixed samples. The technique has a significantly higher throughput and can be used for a wider range of species and more complex samples compared to traditional, non-sequencing techniques. However, mainstream uptake and accessibility of DNA metabarcoding as a general method for biodiversity analysis would benefit from sequencing and data analysis becoming easier to use and cheaper. The MinION sequencer of Oxford Nanopore provides a portable and inexpensive sequencing solution that meets these requirements as it requires minimum lab equipment and provides very quick turnaround of results [1]. The integration of MinION sequencing technology into metabarcoding workflows addresses critical challenges encountered in traditional DNA metabarcoding approaches. For example, the extended read length of Nanopore sequencing enables the resolution of intricate genetic diversity within environmental samples and facilitates the identification of novel or divergent taxa, which may be overlooked by shorter-read technologies.

Metabarcoding data analysis, however, is not yet mainstreamed and many different bioinformatic pipelines have become available over recent years [2,3,4,5,6]. As the sequencing error profile of MinION sequencing differs to that of other sequencing devices such as Illumina sequencers [3,7], analysis pipelines developed with data from other platforms in mind may not perform well on MinION data. Numerous bioinformatic pipelines are available for the commonly used Illumina sequencing (e.g., QIIME2, JAMP, OBITools), but there is still a gap when it comes to metabarcoding data produced through nanopore sequencing. Previously, a pipeline was developed by Voorhuijzen-Harink et al. [2] to identify fish species from DNA-metabarcoding data obtained by MinION sequencing of complex mixtures. This dedicated bioinformatics workflow was developed using freely available tools. At its core, the workflow processes MinION data to produce multiple high-quality consensus DNA barcodes suitable for multi-species identification.

The Voorhuijzen-Harink et al. pipeline was designed to analyze individual samples and does not perform taxonomic comparison between samples. Its throughput is hampered by a lack of compatibility with high-performance computing cluster (HPC) parallelization and the use of an online database to retrieve taxonomy data. The latter furthermore impacts its ability to perform reproducible analyses.

This paper introduces PIMENTA, a PIpeline for MEtabarcoding through Nanopore Technology used for Authentication. PIMENTA is a pipeline for rapid taxonomic identification in samples using MinION metabarcoding sequencing data. This is an updated version of Voorhuijzen-Harink et al. pipeline. We evaluated PIMENTA performance using multiple constructed mock mixtures of commonly encountered zooplankton species of a temperate estuary. Use of mock mixes is recommended because they contain known sequences at a known concentration, which makes it easier to verify the results of the pipeline [8]. The evaluation shows the enhanced detection provided by PIMENTA, which holds promise to improve our comprehension of species-diverse ecological systems.

## 2. Material and Methods

Molecular approaches are increasingly being used to assess the biodiversity of communities of organisms, including zooplankton [9,10]. The diverse group of zooplankton includes many taxa that may have their entire life cycle in the plankton (holoplankton) or only spend part of their lifetime in the water column, including larvae of fish and benthos (meroplankton). The larvae of the latter group are often hard to identify by morphological identification at species or even genus level. Molecular approaches may therefore be especially valuable in ecosystems where larvae and cryptic species contribute substantially to the zooplankton community [11]. Therefore, we chose to evaluate the pipeline on mock samples of zooplankton species.

Multiple adjustments have been made compared to Voorhuijzen-Harink et al., such as parallelization of the analysis of multiple samples with the use of high-performance computing (HPC), the usage of a local taxonomy database and expanded taxonomic results summary.

The pipeline is formed by linking several tools together, resulting in a four-phase data-analysis pipeline:

1) pre-processing: demultiplexing, trimming sequencing adapters, quality trimming, and filtering of the reads,
2) clustering the reads per sample, followed by multiple sequence alignment (MSA) and consensus building per cluster,
3) reclustering consensus sequences from all samples, followed by MSA and consensus building for each cluster,
4) Taxonomic identification.

We benchmark the performance (i.e., the number and taxonomic level of identified zooplankton species) and runtime of PIMENTA with two other existing pipelines, the previous pipeline published by Voorhuijzen-Harink et al.[2] as well as Decona [4]. An overview of the procedures, analysis and databases applied in this study is shown in Figure 1.

**Figure 1:**
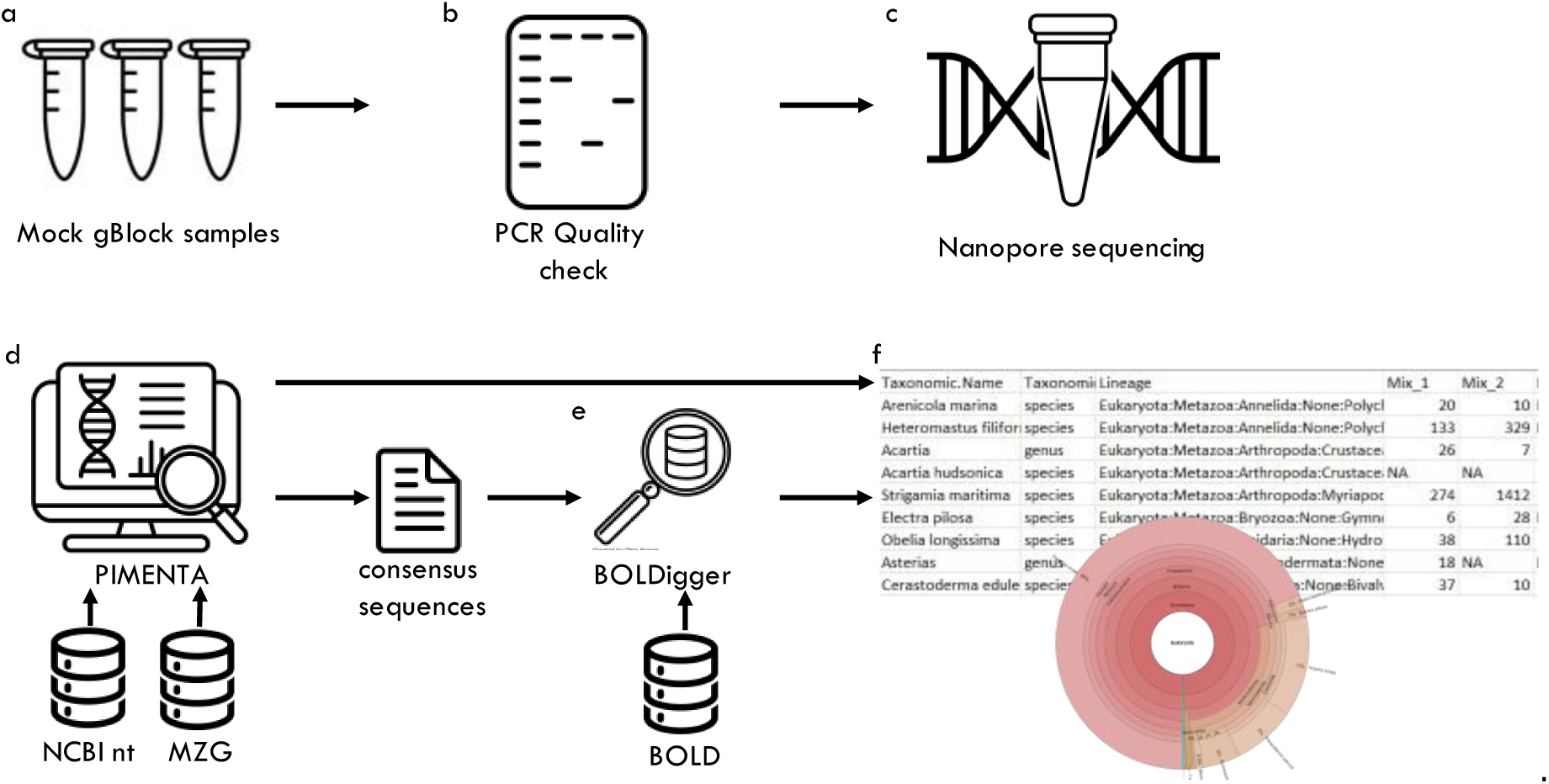
PIMENTA evaluation workflow (A) design synthetic gBlocks sequences using accessions on the NCBI nucleotide database (section 2.1.1); (B) test gBlock sequences for accuracy with PCR (C) sequence gBlock mock mixes with MinION (section 2.1.1); (D) analysis with PIMENTA, using NCBI nt database and MetaZooGene (MZG) database (section 2.1.4, 2.2); (E) BOLDigger analyzes the consensus sequences from PIMENTA with BOLD database (section 2.1.5); (F) Taxonomic overview is created by PIMENTA including (among others) species, taxonomic tree and read count per sample. Krona plots are generated per marker per sample in the PIMENTA pipeline (section 2.2).

### 2.1 Materials and methods

#### 2.1.1 Mock mixes

Fifteen mock mixes were constructed to evaluate the three markers used in this study, as well as the efficacy of the bioinformatic pipelines. Since we had no suitable biological material available on various set of relevant zooplankton species, we chose to use gBlocks^tm^ (IDT): synthetic sequences designed using accessions in the NCBI nucleotide database. gBlocks were selected for species from different taxonomic groups, representing a natural zooplankton community in a temperate estuary. gBlocks were mixed according to table 1, using both equal and skewed proportions. Individual gBlocks were tested using PCR with input of 10^5^ µl-1 copies as described below for the COI, 18SV4 and 18SV9 regions and presence of amplicon was verified with gel electrophoresis. The PCR analysis of the gBlocks was performed to confirm the accuracy of the gBlock sequence. Three of the gBlocks did not produce an amplicon and were excluded from the evaluation: *Semibalanus balanoides* for COI and *Clupea harengus* and *Strigamia maritima* for 18SV4.

**Table 1.** Mock mixes (MM) distribution. Values are given in fractions.

| Species | Common Name | Phylum | Mix |  |  |  |  |
| --- | --- | --- | --- | --- | --- | --- | --- |
|  |  |  | 1 | 2 | 3 | 4 | 5 |
| *Heteromastus filiformis* | Red Thread Worm | Annelida | 0.10 | 0.20 | 0.01 | 0.001 | 0.11 |
| *Arenicola marina* | European lugworm | Annelida | 0.10 | 0.03 | 0.01 | 0.001 | 0.11 |
| *Acartia tonsa* |  | Arthropoda | 0.10 | 0.03 | 0.91 | 0.001 | 0.11 |
| *Semibalanus balanoides* | Common barnacle | Arthropoda | 0.10 | 0.03 | 0.01 | 0.001 | 0.11 |
| *Strigamia maritima* |  | Arthropoda | 0.10 | 0.03 | 0.01 | 0.001 | 0.11 |
| *Electra pilosa* | Hairy Sea-mat | Bryozoa | 0.10 | 0.20 | 0.01 | 0.001 | 0.11 |
| *Clupea harengus* | Atlantic herring | Chordata | 0.10 | 0.20 | 0.01 | 0.001 | 0.11 |
| *Obelia longissima* | Sea Fur | Cnidaria | 0.10 | 0.20 | 0.01 | 0.001 | 0.11 |
| *Asterias rubens* | Common starfish | Echinodermata | 0.10 | 0.03 | 0.01 | 0.001 | 0.01 |
| *Cerastoderma edule* | Common cockle | Mollusca | 0.10 | 0.03 | 0.01 | 0.991 | 0.11 |
| sum |  |  | 1.00 | 1.00 | 1.00 | 1.00 | 1.00 |

All primers were synthesized (Biolegio, NL) with the standard Oxford Nanopore adapters (Oxford Nanopore Technologies, UK) 5’-TTTCTGTTGGTGCTGATATTGC-3’ for the forward primers and 5’-ACTTGCCTGTCGCTCTATCTTC-3’ for reverse primers. Each PCR reaction contained 1x PCR Buffer, 2.5 mM MgCl2, 200 μM dNTPs, 250 nM of each primer, 5% trehalose, 1.25 μg BSA, 1 U Platinum™ Taq DNA Polymerase (Invitrogen, USA), 4 ng template DNA and nuclease-free water to an end-volume of 25 µl. A “touchdown” PCR program was used in a Veriti Thermo Cycler (Thermo Fisher Scientific, USA) for all primer pairs, starting with initial denaturation for 2 min at 94°C, followed by 15 cycles: denaturation for 30 sec at 94°C, annealing for 3 min at 56°C (touchdown of −1°C per cycle) and extension for 60 sec at 72°C, followed by 20 cycles: denaturation for 30 sec at 94°C, annealing for 3 min at 42°C temperature and extension for 60 sec at 72°C and last a final extension of 10 min at 72°C. Two replicate PCRs were performed for each mock mix sample in separate PCR runs, and were pooled before sequencing. Water controls were included for all runs. Amplicons were checked with gel electrophoresis and stored at –20 °C.

#### 2.1.2 DNA extraction

To remove the ethanol added to samples for preservation and storage purposes, 1/33 volume of Sodium acetate (pH 5.2, 3M) was added to each sample to remove the ethanol and to precipitate the DNA. Samples were centrifuged for 60 min at 3800x g at room temperature, supernatant discarded, and pellets were collected for DNA extraction. DNA extraction was performed with the Qiagen DNeasy Blood and Tissue Kit (Qiagen) according to the manufacturer’s protocol with the following adjustments: the volume of buffer ATL and Proteinase K were adjusted in ratio to the amount of material. Lysis was performed overnight at 56°C while shaking. 200 µl of the lysate was taken and processed according to manufacturer’s protocol. Elution was done with 200 µl buffer AE.

#### 2.1.3 MinION library prep & sequencing

8 µl of PCR product of each mock mix per marker was combined and subsequently purified using the Qiagen QIAquick Purification kit (Qiagen, Germany) according to the manual. The DNA was quantified using the Quantus Fluorometer (Promega, United States), and diluted to 10 ng/µl. Per sample, 1 µl was combined with 1.5 µl of Barcode Primer (Oxford nanopore, England) and 25 µl of LongAMP Hot Start Taq 2x MM (New England Biolabs, United States) and subsequently mixed gently. This mixture was incubated at 95°C for 3 min, followed by 15 cycles of 95°C for 15 sec, 56°C for 15 sec and 65°C for 60 sec using a thermal cycle, and a final extension at 65°C for 6 min after which samples were cooled down to 8°C until further use. The resulting library was washed by adding 40 µl of pre-vortexed AMPure xp beads (Beckman Coulter, United States) and rotated by hand for 5 min at room temperature, spun down and separated using a DynaMag-96 magnetic rack (Invitrogen, United States) for 5 min. Once the eluate was clear the supernatant was pipetted off and washed twice with freshly prepared 70% (v/v) ethanol. The washed pellet was dried briefly and resuspended in 10 µl of 10 mM Tris-HCL, 50 mM NaCl with a pH of 8.0. After incubating in the magnetic rack for 2 min at room temperature, the eluant containing the library was transferred to a clean 1.5 ml LoBind tube. Again DNA was quantified using the Quantus, after which to a maximum of 5 barcoded libraries were pooled, and diluted to a total of 30 ng DNA (evenly distributed over n libraries) in 4 µl. 1 µl of rapid adapter (Oxford Nanopore Technologies, England) was added to the mixture, gently mixed, spun down and incubated for 5 min. at room temperature and subsequently prepared for loading onto a FLO-FLG001 Flongle flow cell (Oxford Nanopore Technologies, England) according to the manufacturer’s manual. In short, the flow cell was primed by opening the sample port and loading 120 µl of priming mix (3 µl Flush Tether to 117 µl of Flush Buffer) and 30 µl of sample mix (15 µl of Sequencing Buffer, 10 µl pre-mixed loading beads and 5 µl of DNA library) into the flow cell via the sample port ensuring no air bubbles were introduced. The sample port was closed, and sequencing was started using MinKNOW software (version 22.05.5). Sequencing was stopped after 30000 – 50000 reads per sample.

#### 2.1.4 Databases

The NCBI nt database, updated in May 2022, was used to create the main database used in PIMENTA. The Barcode of Life Data System (BOLD) [16] database has been utilized in October 2022, using its COI barcode annotation database. The 18S and COI databases from the MetaZooGene Database (MZGdb) [17] were downloaded in October 2022.

#### 2.1.5 BOLDigger

BOLDigger [18], a tool that streamlines the process of querying DNA sequences against the BOLD database, was used to analyze COI consensus sequences produced by PIMENTA to see if the results were equivalent to utilizing the NCBI nt database.

### 2.2 Bioinformatic analysis

#### 2.2.1 PIMENTA data processing workflow

PIMENTA consists of four subsequent phases: preprocessing, analysis per sample, all samples combined, and taxonomic identification. The pipeline with instructions can be found on GitHub: https://github.com/WFSRDataScience/PIMENTA

For this study, the following settings were employed; however, users can utilize their preferred settings.

##### Preprocessing

Basecalling and demultiplexing of the raw FAST5 files is performed using Guppy GPU version 6.1.1 keeping only FASTQ files [19]. Sample specific barcodes are trimmed by Guppy as well.

##### Analysis per sample

FASTQ reads with an average Phred score of at least 12 and read length between 10 and 1100 bases are selected using PRINSEQ 0.20.4 [20]. Clustering is performed separately for each sample using cd-hit-est 4.6.7 with 93% identity with a minimum cluster size of one [21]. Multiple sequence alignment (MSA) is performed on the sequences within each cluster using MAFFT v7.471 with adjust direction enabled and with the L-INS-i iterative refinement method selected[22], L-INS-i can align a set of sequences containing sequences flanking around one alignable domain. For clusters with more than 100 reads, 100 reads are randomly selected prior to MSA. Consensus sequences are generated from the MSA by selecting nucleotide positions with a percent identity exceeding 30%. These positions represent the most frequently occurring nucleotides in the alignment.

##### Analysis of all samples combined

The consensus sequences from all individual samples are combined and then reclustered using cd-hit-est with 100% identity. MSA is performed using MAFFT, and consensus sequences are extracted with the same settings. Sequences are separated into pools representing the different markers (COI, 18SV4 or 18SV9) by detecting the DNA barcode primer sequences used to amplify these regions using Cutadapt version 4.2[23]. Barcode primer sequences are trimmed with a 15% maximum acceptable error rate and a minimum overlap of 25 nt. To assign consensus sequences to taxonomy, standalone BLASTn megablast searches (BLAST+ v2.12.0) [24] are performed against the NCBI (National Centre for Biotechnology Information) nt database (accessed May 2022). Hits with ≥ 90% sequence identity and ≥ 90% query coverage are further considered. A list of accessions and taxa (environmental samples etc.) are removed from the BLAST output. See “2.2.2 Filtering BLAST database and taxonomy” for more information about how the BLAST output was assessed for taxonomic identification.

#### 2.2.2 Filtering BLAST database and taxonomy

The BLAST results were filtered as follows. A manually curated list (Additional File 1) was created to filter out taxa, and another list to filter out specific sequences (Additional File 2). The taxa that were removed are predominantly higher-level taxa and unplaced species and genera that appeared in database entries based on environmental samples. Taxa without a family or genus were only included in the results if there were no other taxa within the same family or order with a marker (e.g., COI) sequence in the database. The list of excluded sequences comprises sequences that could lead to false positive assignments as they were annotated to be derived from species from another family or order than that of all other similar sequences in the database. Briefly, when processing BLAST results if a sequence had a species annotation that was distinct from the rest of the results and this species annotation belonged to a different family or order than the other results, an additional check were performed. To check, the sequence was blasted against the NCBI nt database. If again all results were from a different family/order with a high identity and coverage (>98%), the sequence was added to the list. This procedure is expanded from the pipeline used by Voorhuijzen-Harink et al, which does not perform filtering of taxa.

Another filtering step was applied by only selecting results within the kingdom Metazoa, which removes all bacteria/fungi/plants from the taxonomy output. This additional filter is adjustable in the settings file, making it possible to analyze several types of data.

The BLAST output was interpreted following these guidelines: all hits with the top bit score were selected. If there were multiple hits with the same score and subject sequences were annotated as belonging to different assigned species, the cluster fragment was considered to lack the discriminatory power to refer the hit to species level. In such cases, the sequence would then be downgraded to a genus-level identification. Similarly, if the hits have different assigned genera, the cluster fragment was then downgraded to a family-level identification.

#### 2.2.3 Evaluation

PIMENTA is evaluated on three criteria: 1) read usage, 2) taxonomic identification, and 3) runtime of the pipeline. Samples were analyzed in parallel on a high-performance computing cluster (HPC) with an assignment of 15 CPU cores and 60 GB total RAM per sample.

We benchmarked PIMENTA against two other pipelines: Voorhuijzen-Harink et al. pipeline [2] and Decona 1.3.1 (https://github.com/Saskia-Oosterbroek/decona). For the comparison to Decona, default settings were used except for the following alterations in table 3.

**Table 2.**
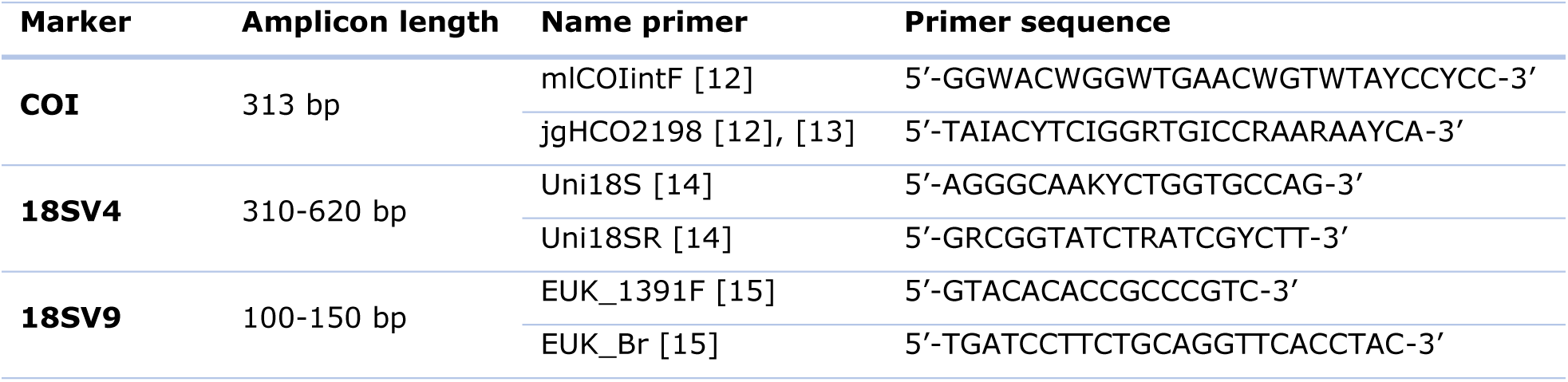
marker primers.

**Table 3:**
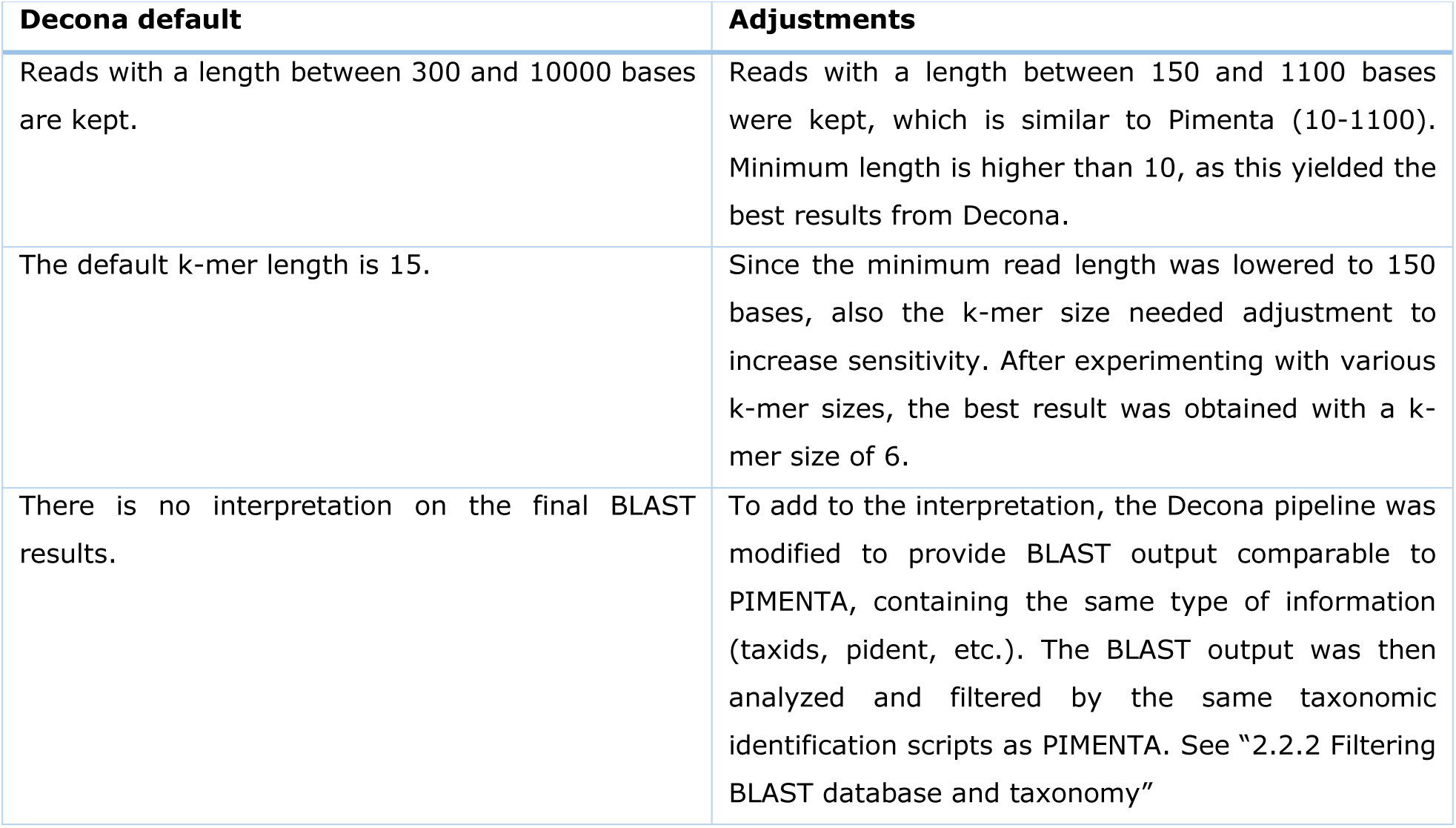
Adjustments to the Decona pipeline, to make the results comparable to PIMENTA.

## 3. Results and discussion

This study demonstrates the capabilities of the improved nanopore DNA metabarcoding pipeline PIMENTA. The addition of a reclustering step and tweaking of other steps, for example clustering identity, has improved accuracy and reduced runtime over the previous implementation. Although our pipeline has been successful, there are still some challenges in DNA metabarcoding.

### 3.1 Rework and expansion of the Voorhuijzen-Harink pipeline

As previously stated, PIMENTA is an updated version of the Voorhuijzen-Harink et al. pipeline. The key difference is the inclusion of reclustering all samples together, to compare samples more effectively. Barcode trimming is now done by Guppy instead of Porechop. Because MinION sequencing produces to date higher quality reads compared to 2019 [2], clustering is now done at 93% rather than at 80% identity. An additional filter has been implemented to exclude BLAST hits to sequences annotated at high taxonomic rank for which other hits to an identical sequence with a more specific taxonomic identification were also found. The sequences that are removed are often annotated as being derived from environmental samples, amongst similar terms.

Another notable enhancement is the capability to initiate the pipeline at any specific stage, simplifying the process of examining data using various configurations without the need to re-execute the entire process. Additionally, the pipeline’s installation process has been streamlined, eliminating the requirement to separately install all tools and packages other than Guppy.

The taxonomic identification script is modified to make it faster and less error prone; the previous version was written in R, and the current version is written in Python. The R script needs a connection to an online database, which decreases the speed, while the Python script uses a local database. This also improves reproducibility, with more version control compared to an online database.

Krona plots are also included to provide a visual representation of the results. The output of the taxonomic table is enhanced to incorporate more information, such as taxonomic trees.

### 3.2 Read utilization in analyses

PIMENTA consistently provided taxonomic identification for more reads than Decona across all mixes (T-test P < 0.05, df = 6.57, t = 15.92; Figure 3). Decona provides fewer identifications most likely due to its approach to filtering database matches. Though the difference is non-significant (P = 0.75, df = 7.68, t = –0.33), the Voorhuijzen-Harink et al. pipeline provided taxonomic identification for slightly more reads overall than PIMENTA. This is explained by the filtering of taxids that was added to PIMENTA, which removes some ultimately less informative matches to the database.

**Figure 2:**
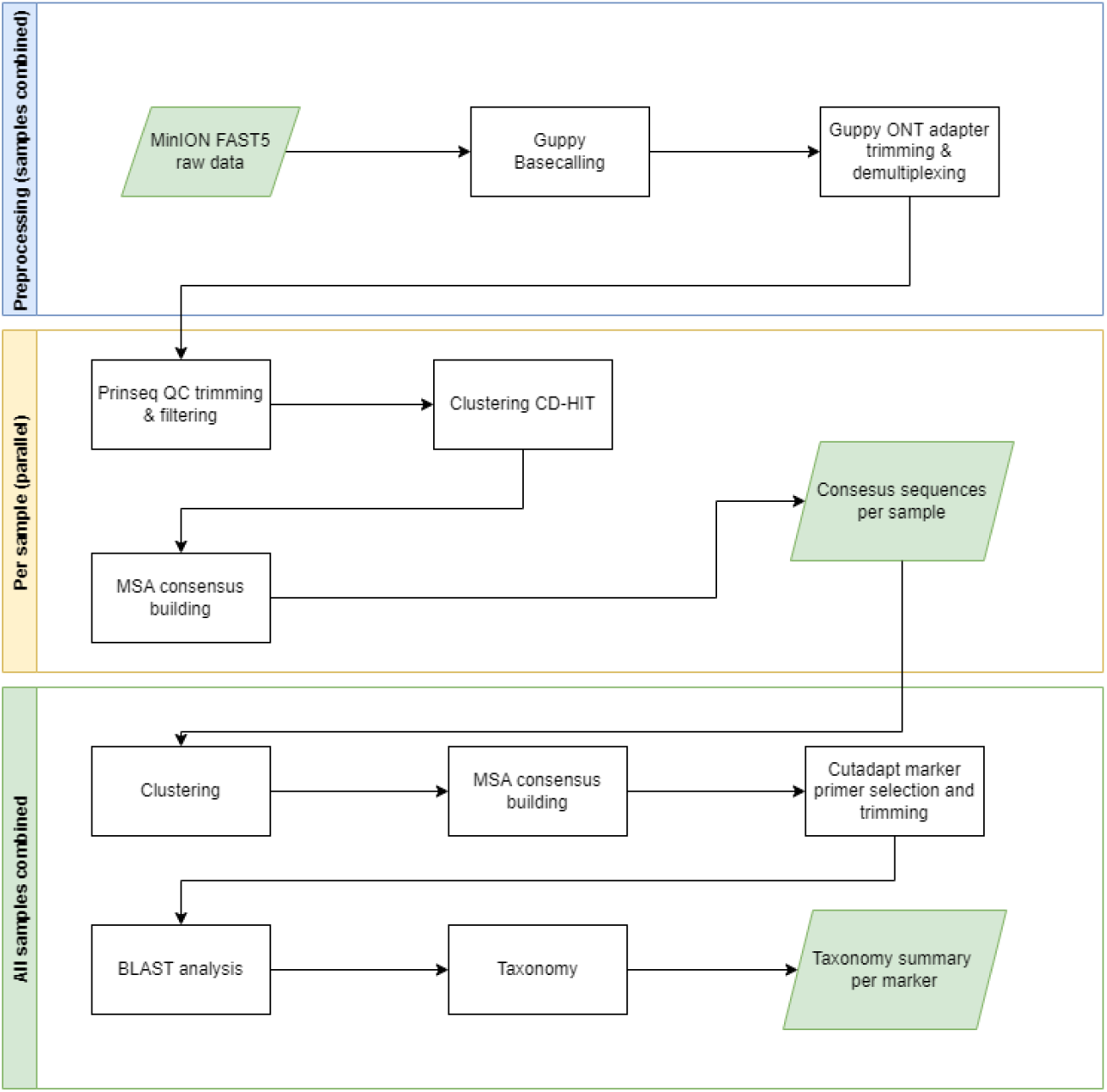
Workflow of PIMENTA. PIMENTA consists of three different sections: Preprocessing, analysis per sample, all samples combined. Preprocessing consists of basecalling, demultiplexing and adapter trimming. The next section “Per sample” analyses all samples in parallel and returns a file with consensus sequences per sample. With the last section, “all samples combined”, the workflow identifies the taxonomy for all samples combined.

**Figure 3:**
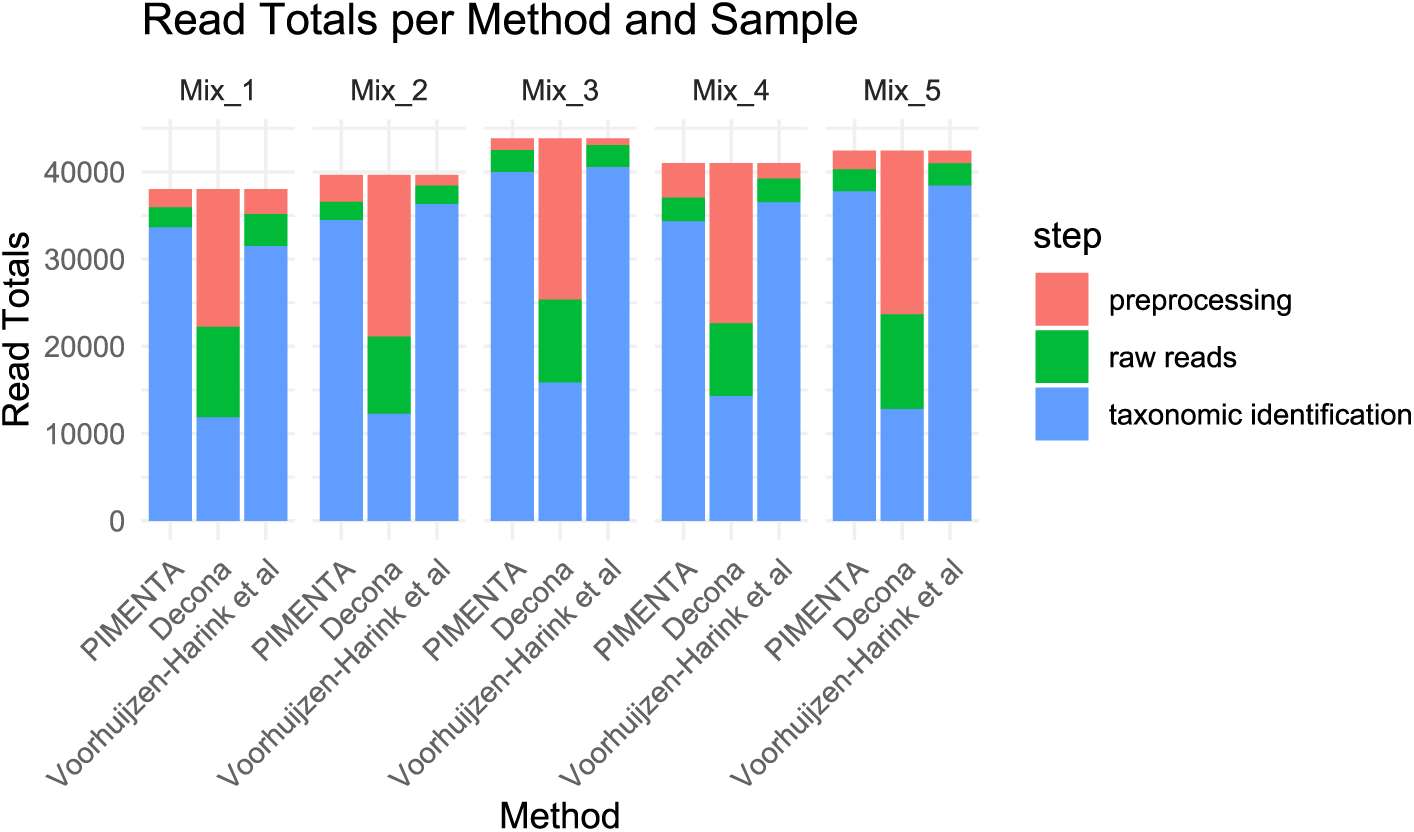
Distribution of reads at various stages of data processing. The uppermost segment of the bars corresponds to the initial dataset, comprising all the raw reads (red). The middle segment represents the preprocessing phase (green), which includes the reads marked as taxonomic identification as well as a subset of reads that have undergone preliminary processing steps, such as quality filtering and adapter removal. The bottom segment (blue) of the figure comprises reads that have successfully received taxonomic identification.

Figure 4 shows differences between markers and tools in terms of the numbers of reads that were taxonomically classified, highlighting the strengths and weaknesses of each pipeline when analyzing the different markers across different samples. 18SV9 consistently exhibits the highest assigned read counts across all samples, suggesting its prevalence or ease of detection. By contrast, COI and 18SV4 have lower read counts.

**Figure 4:**
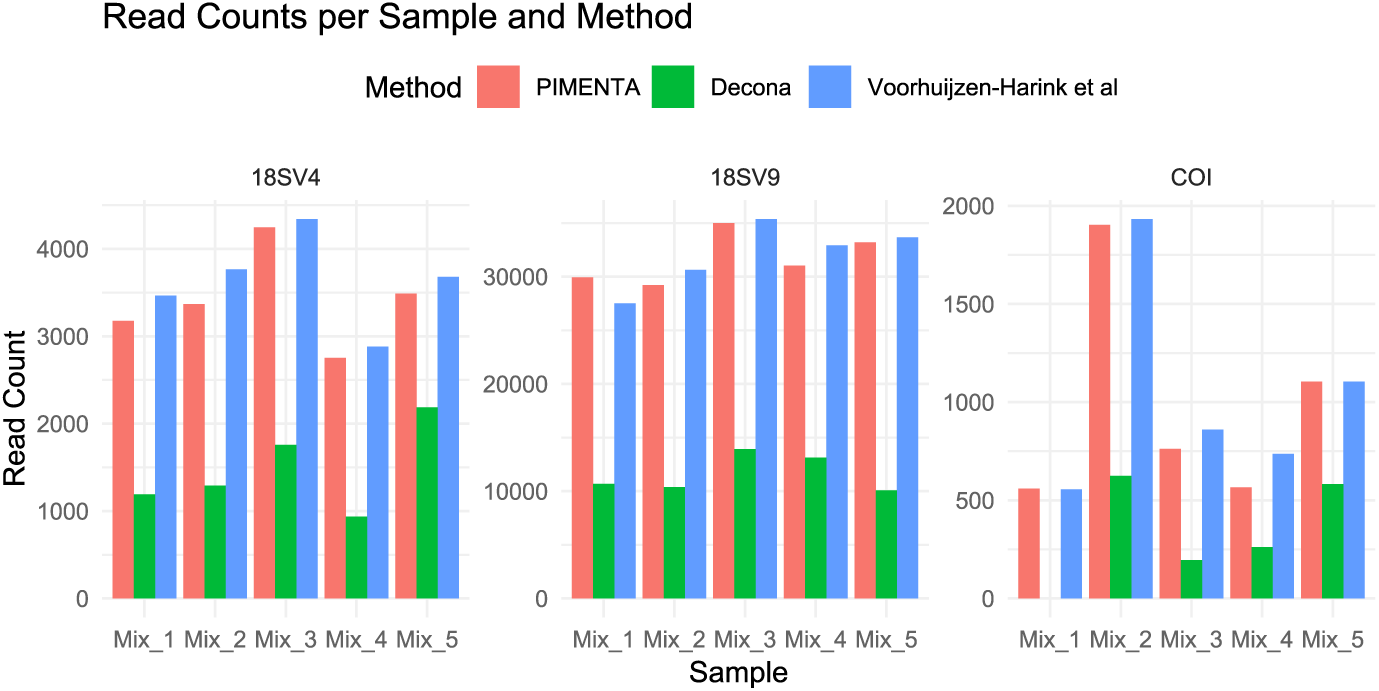
Total reads used for taxonomic identification with PIMENTA, Decona and Voorhuijzen-Harink et al pipeline. These reads have undergone a taxonomic identification process via BLAST and BLAST filtering, and the displayed counts exclude any reads that were filtered out during earlier stages of data processing. It showcases the number of reads that successfully navigated through the pipelines, leading to taxonomic assignments. Note the different ranges on the Y-axis.

### 3.3 DNA metabarcoding sensitivity and specificity

The ideal metabarcoding experiment is capable of detecting traces of DNA at low relative abundance and accurately identifying the species from which it originates. We assessed the performance of the method including the pipelines using mixes of known relative abundance of synthesized DNA representing different species in the form of gBlocks. The presence of all gBlocks was reliably identified based on at least one marker by at least one pipeline, with the exception of the gBlock representing *Clupea harengus*. As there were no or extremely few reads assigned to this species by all pipelines for all markers we have omitted this species from the results. PIMENTA assigned reads from most gBlocks successfully at species level and was able to detect all gBlocks at family level in equal or smaller relative abundance (Table 4). The taxonomic rank at which a sequence can be identified differed between markers and pipelines, with PIMENTA being capable of detecting most included gBlocks at species level across all markers (Table 5).

**Table 4:**
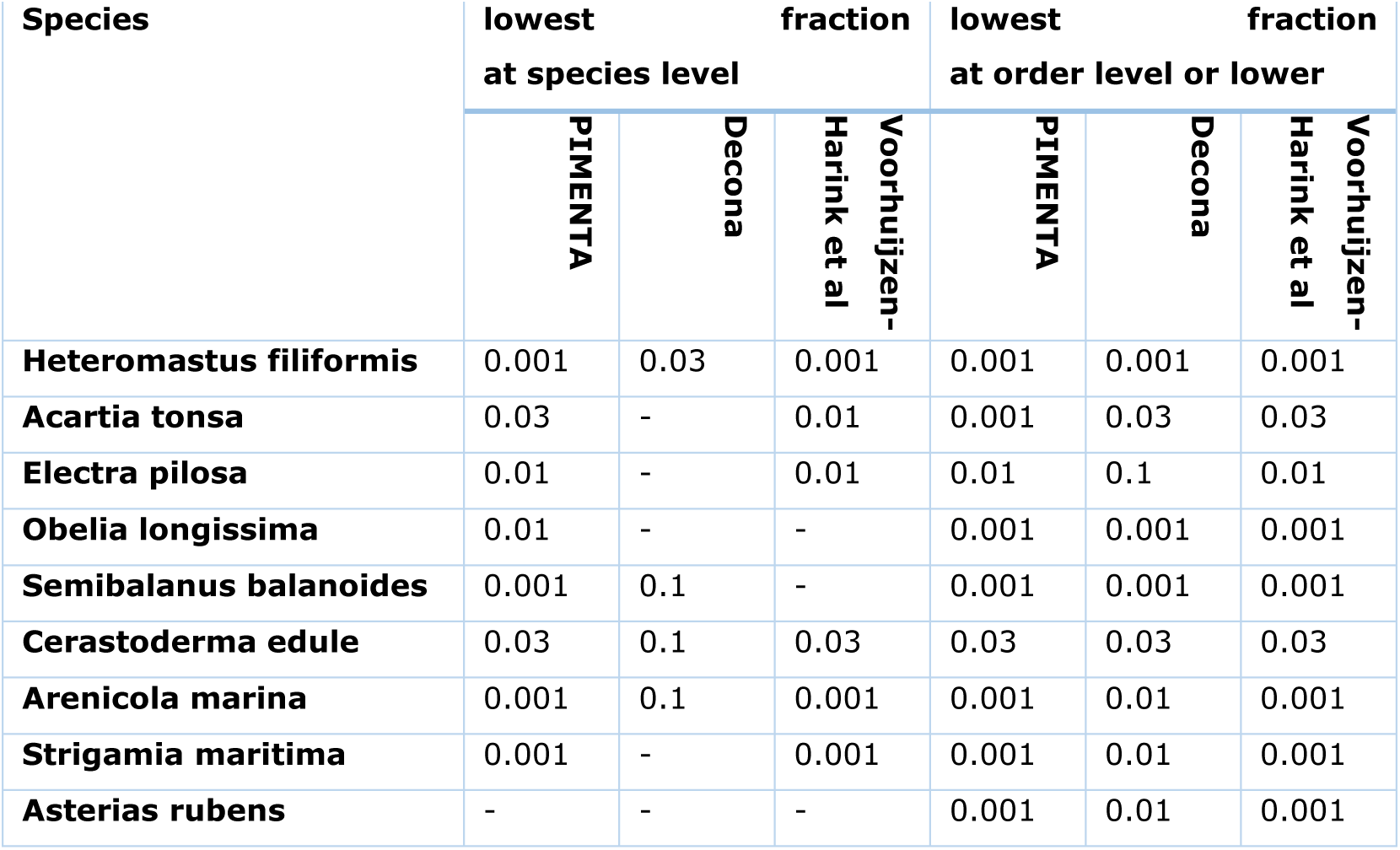
Lowest fractional abundance found by PIMENTA, Decona and Voorhuijzen-Harink et al at species level and at order level or lower taxonomic rank.

| Species | lowest fraction at species level |  |  | lowest fraction at order level or lower |  |  |
| --- | --- | --- | --- | --- | --- | --- |
|  | PIMENTA | Decona | Voorhuijzen-Harink et al | PIMENTA | Decona | Voorhuijzen-Harink et al |
| **Heteromastus filiformis** | 0.001 | 0.03 | 0.001 | 0.001 | 0.001 | 0.001 |
| **Acartia tonsa** | 0.03 | - | 0.01 | 0.001 | 0.03 | 0.03 |
| **Electra pilosa** | 0.01 | - | 0.01 | 0.01 | 0.1 | 0.01 |
| **Obelia longissima** | 0.01 | - | - | 0.001 | 0.001 | 0.001 |
| **Semibalanus balanoides** | 0.001 | 0.1 | - | 0.001 | 0.001 | 0.001 |
| **Cerastoderma edule** | 0.03 | 0.1 | 0.03 | 0.03 | 0.03 | 0.03 |
| **Arenicola marina** | 0.001 | 0.1 | 0.001 | 0.001 | 0.01 | 0.001 |
| **Strigamia maritima** | 0.001 | - | 0.001 | 0.001 | 0.01 | 0.001 |
| **Asterias rubens** | - | - | - | 0.001 | 0.01 | 0.001 |

**Table 5:** Lowest taxonomic rank found by PIMENTA (P), Decona (D) and **Voorhuijzen-Harink et al** pipeline (V) per species with markers COI, 18SV4 and 18SV9. The mixes do not contain all markers of each species, the excluded markers are specified with an asterisk. S=species, G=genus, F=family, C=class, O=order,”-” indicates the species is missing from the results.

| Species | marker |  |  |  |  |  |  |  |  |  |  |  |
| --- | --- | --- | --- | --- | --- | --- | --- | --- | --- | --- | --- | --- |
|  | Any |  |  | COI |  |  | 18SV4 |  |  | 18SV9 |  |  |
|  | PIMENTA | Decona | Voorhuijzen-Harink et al | PIMENTA | Decona | Voorhuijzen-Harink et al | PIMENTA | Decona | Voorhuijzen-Harink et al | PIMENTA | Decona | Voorhuijzen-Harink et al |
| *Heteromastus filiformis* | S | F | S | S | S | S | S | S | S | S | F | S |
| *Acartia tonsa* | S | F | S | G | F | G | S | F | S | S | F | S |
| *Electra pilosa* | S | O | S | S | - | S | O | O | O | G | - | - |
| *Obelia longissima* | S | G | G | S | F | G | G | G | G | G | G | G |
| *Semibalanus balanoides* | S | F | O | * | * | * | S | S | O | O | F | O |
| *Cerastoderma edule* | S | S | S | S | S | S | S | S | S | S | S | S |
| *Arenicola marina* | S | F | S | S | - | S | S | S | S | S | S | F |
| *Strigamia maritima* | S | - | S | S | F | S | * | * | * | S | F | S |
| *Asterias rubens* | G | F | G | G | - | G | G | - | G | O | F | O |

The lowest taxonomic ranks identified by Decona varied more compared to PIMENTA. Some species were consistently identified at species level, while others were identified at the family or order level. The two methods also differ in their ability to identify certain species. PIMENTA was able to identify *Electra pilosa*, *Obelia longissima* and *Strigamia Maritima* at species level in at 0.1 relative abundance. Decona was not able to provide the same level of resolution for these species. The main differences between Decona and PIMENTA that could underlie the difference in outcome is that Decona blasts the complete consensus sequence, whereas PIMENTA extracts the marker segment from each consensus sequence to get more distinct matches, and that PIMENTA uses a higher clustering threshold. The Voorhuijzen-Harink et al. pipeline produces results more similar to those of PIMENTA, but PIMENTA detected species that were not detected by Voorhuijzen-Harink et al. and was able to identify more gBlocks down to species levels, such as *Semibalanus balanoides* at 0.001 abundance. This is likely due to a higher clustering identity and added filtering of BLAST results, resulting in a more accurate identification.

The resolution of identification also differs between markers. Only COI allowed for identification of *O. longissima* at species level; based on reads of 18S4 and 18SV9 identification was only accurate to genus level. COI has a low read count in general and showed and the fewest species-level assignments but has the highest resolution out of the three markers. Yield of the COI marker can potentially be improved by treating markers as individual samples rather than pooling them together for each biological sample. Overall, species were identified at lower taxonomic rank if the COI primers were able to amplify the species-specific COI sequence.

Generally, PIMENTA found the most species in each sample with each marker and had the lowest number of misidentifications. An overview of the relative abundance assigned to each species is provided in Figure. Compared to the Voorhuijzen-Harink et al. pipeline, PIMENTA included fewer reads that were assigned to a taxonomic rank higher than order. Voorhuijzen-Harink et al. identified relatively more reads at kingdom level (Figure 6), which resulted into a higher number of reads classified as other in Figure 5. Though PIMENTA assigns taxonomic information to slightly fewer reads in general, this difference is also due to the removal of non-specifc taxids from the database used by PIMENTA. Decona failed to produce results for Mix_1 with COI and did not identified fewer species based on 18SV4 and COI reads. Using 18SV9 reads, however, Decona outperformed the Voorhuijzen-Harink et al. pipeline, as fewer higher rank classifications / misclassification were provided and more species were found. In the analysis of the sequences of the relatively short markers 18SV4, 18SV9 and COI, PIMENTA outperformed Decona in several aspects. There is an important difference in design consideration between PIMENTA and Decona; whereas PIMENTA is designed with these short markers in mind, Decona was created for the analysis of longer reads. Decona may therefore perform significantly better in benchmarks involving longer markers. More details can be found in Additional File 3.

**Figure 5:**
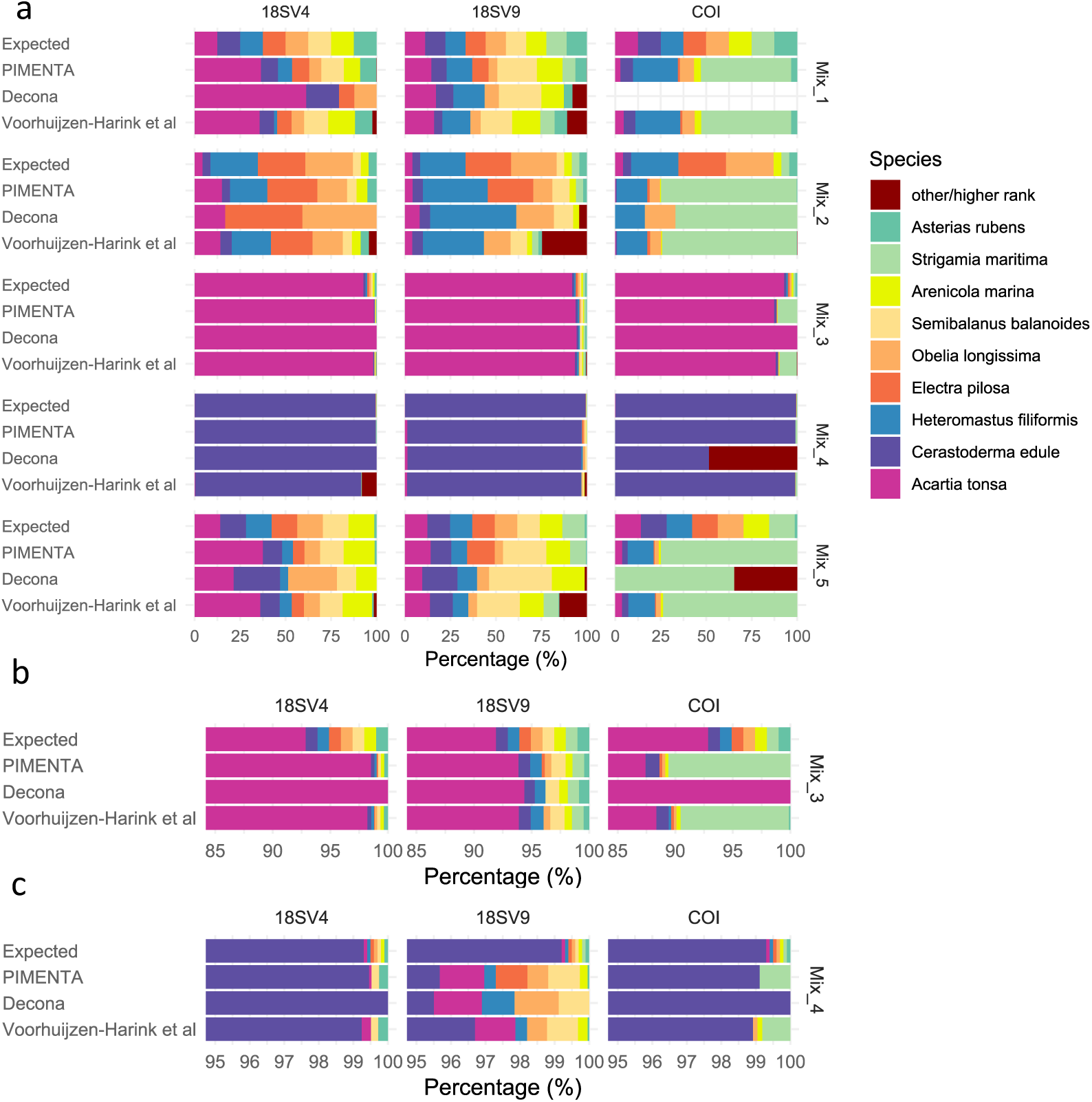
Relative abundances of species included in gBlocks mixes. a. Expected distribution of read attributions compared to the distributions determined by running the pipelines PIMENTA, Decona and Voorhuijzen-Harink et al. Taxonomic identification of the reads ranges from species to order. “Other/higher rank” (in dark red) includes every false positive species and taxonomic identification higher than order. b. Close up to the last 15% of mix_3. c. Close up on the last 5% of mix_4, excluding “Other/higher rank”.

**Figure 6:**
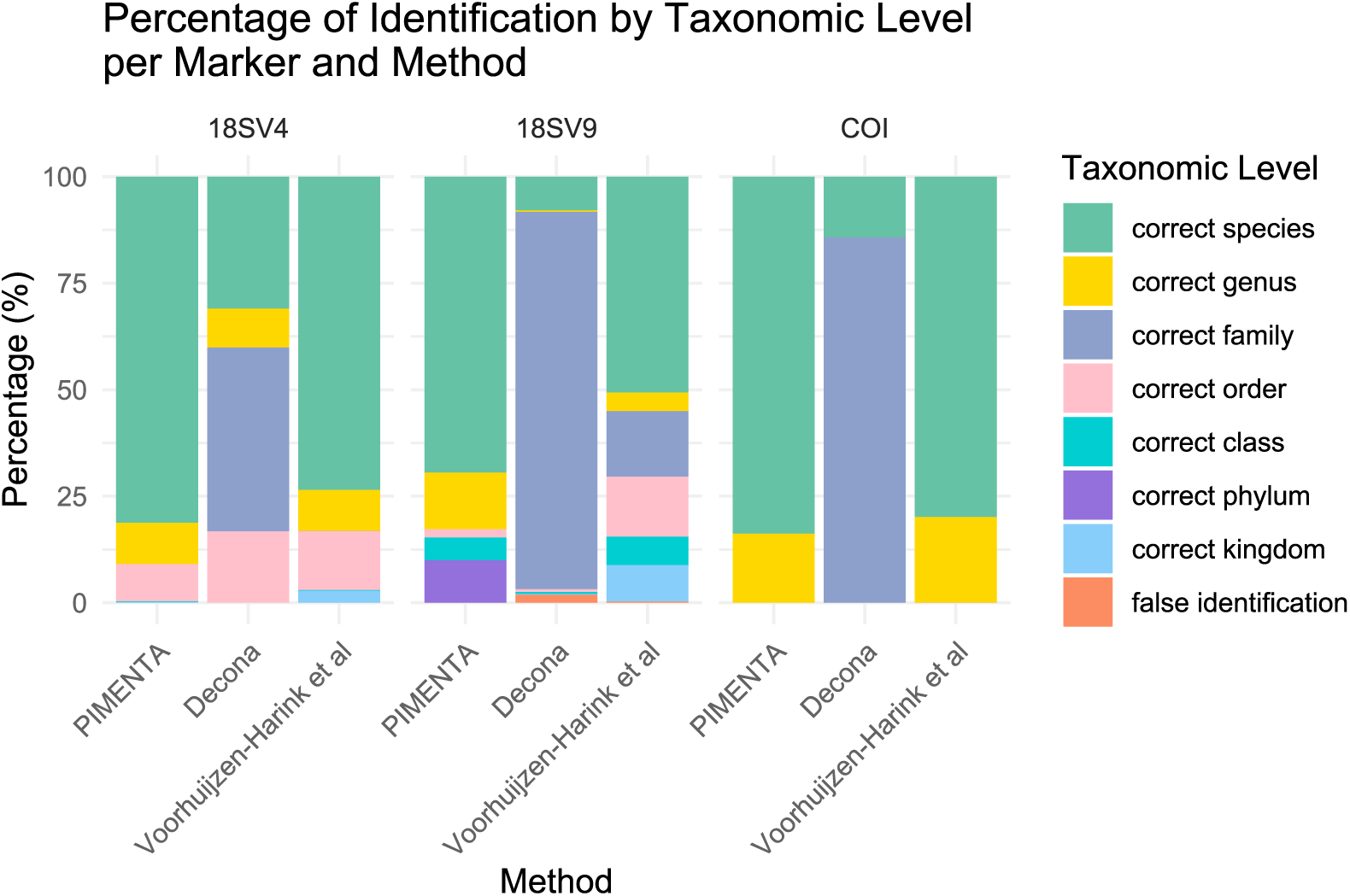
Distribution of read attributions to different taxonomic levels determined by running the pipelines PIMENTA, Decona and Voorhuijzen-Harink et al. The figure allows for a comparative examination of how each pipeline performs in terms of taxonomic assignments across different taxonomic levels, shedding light on the effectiveness of these methodologies in taxonomic identification.

### 3.4 Runtime

The runtime of different modes, parallel and consecutively, were compared using the gBlock mock mix samples (Figure 7). PIMENTA allows analysis steps to be run for each sample in parallel, including quality control (QC) filtering, the first clustering, MSA and consensus building. Decona and Voorhuijzen-Harink et al. have no parallel version. Among the different methods tested, PIMENTA in parallel mode, with a runtime of 45 minutes for analysis of all five gBlock mixes, was the fastest overall. Seventy percent of this time was spent on processing steps which can be performed in parallel. When comparing non-parallel run modes, PIMENTA was still the fastest, taking 90 minutes, followed by Decona at 281 minutes and finally the Voorhuijzen-Harink et al pipeline, which required 292 minutes to complete the analysis.

**Figure 7:**
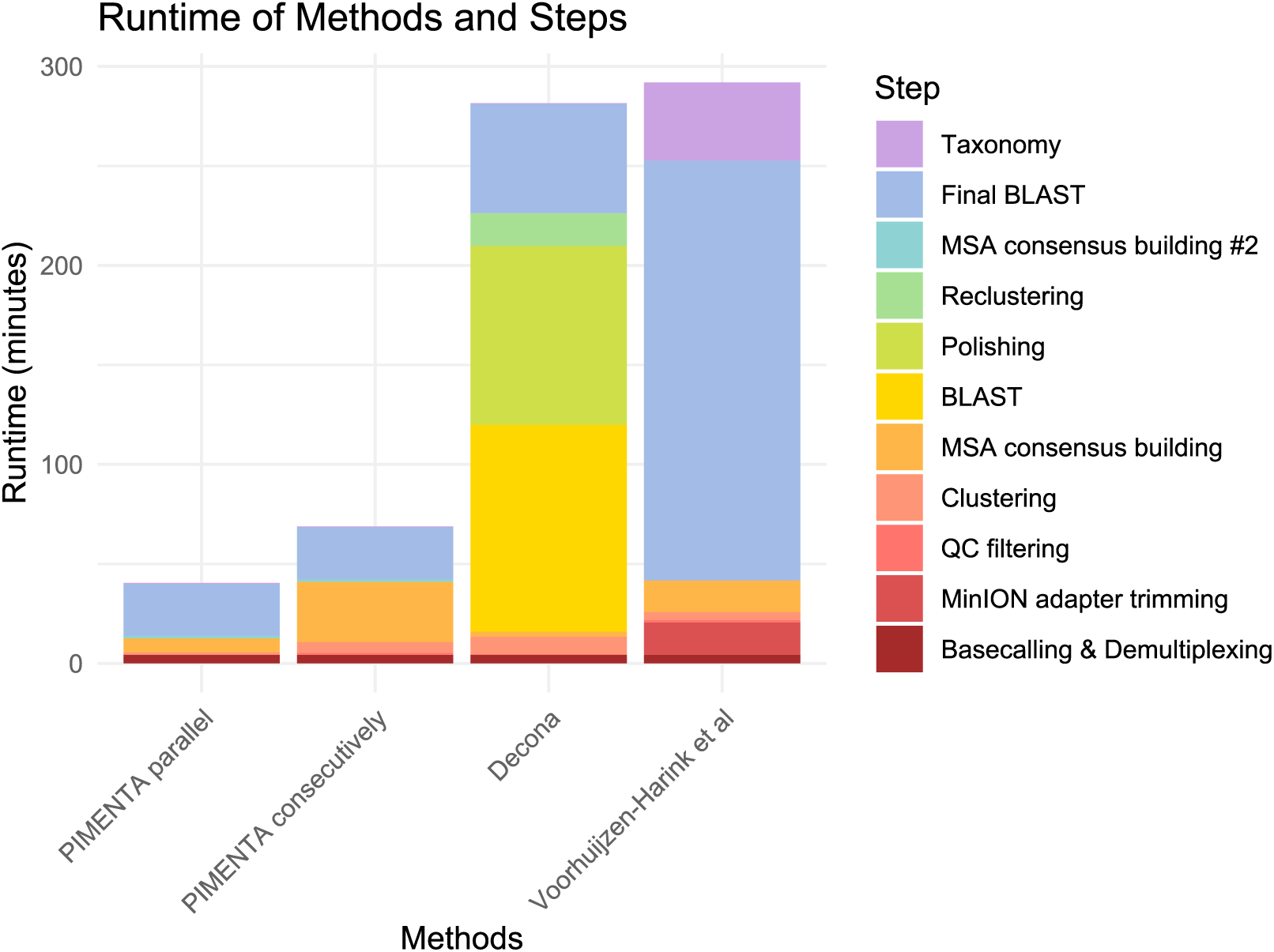
Runtime comparison between PIMENTA, Decona and Voorhuijzen-Harink et al. The runtimes of PIMENTA were determined for both parallel and consecutive run mode.

The differences in run time are due to differences in design of the pipelines. There is a large variation in the time spent on BLAST, which Decona performs twice. Other factors that influence the time taken for BLAST is the number of sequences used as query – e.g., Decona does not recluster samples together, but processes them individually, which often means performing redundant searches. Decona also polishes reads, whereas the other pipelines rely on trimming, Q-score filtering and consensus building to filter out sequencing errors.

### 3.5 Reference databases

Taxonomic identification of marker sequences requires an accurately annotated database with a good representation of different species. In practice, this still a major bottleneck in metabarcoding is the presence of many erroneous sequences and annotations in reference databases that are required for taxonomic classification of the sequences, especially larger general-purpose databases such as NCBI GenBank and nt [10,26,27]. A lot of potentially relevant data has undergone no or limited curation, and the bulk of sequences in the nt database is not informative for metabarcoding analysis. These features increase the time required to process data and potentially lead to incorrect, or inaccurate identifications.

Use of curated marker database, such as MZGdb or BOLD, is one solution. MZGdb [17] is a curated zooplankton database that includes marker sequences from both NCBI and BOLD. The disadvantages of these databases are that the coverage of zooplankton species is still limited – presenting a quality/quantity trade-off – and that the databases are not directly compatible with the different pipelines. To address the latter, we include scripts to utilize MZGdb with PIMENTA for future use when the database coverage has increased. An overview and more information about zooplankton representation in the NCBI nt database and MZGdb can be found in the Additional File 4. For research questions focussed on organisms other than zooplankton, using other curated databases instead of general-purpose databases may improve the quality of the results depending on the organisms of interest.

For matching consensus sequences to entries in the BOLD database, the tool BOLDigger[18] can be used instead of BLAST. We tried BOLDigger on PIMENTA-generated COI clusters in this study and found comparable identifications to those based on BLAST (Additional File 5). A potential downside is that queried sequences are uploaded to the BOLD database, which may not be desired if it is sensitive data. The lack of offline availability further impacts reproducibility. Whether these limitations are acceptable is a decision that will differ between different users.

## Conclusion

PIMENTA was able to achieve rapid and accurate species identification using MinION sequencing while enabling the comparison of multiple samples. PIMENTA provided an improvement in taxonomic accuracy, read usage, and runtime compared to existing pipelines.

## Declarations

**Ethics approval and consent to participate**

Not applicable

## Consent for publication

Not applicable

## Availability of data and materials

PIMENTA is available with instructions and test data at: github.com/PIMENTA-pipeline

## Acknowledgement

We would like to thank Arno Bak, Gijs Kleter, Martijn Staats, Stefan Aanstoot, Viola Kurm for their contributions to this project.

## Supplemental information

Additional File 1:

List with excluded taxids

Additional File 2:

List with excluded NCBI sequence accession ids.

Additional File 3:

Excel file containing taxonomic identification per method

Additional File 4:

Excel file with representation zooplankton in databases

Additional File 5:

Excel file with BOLDigger results PIMENTA COI clusters

